# When sexual selection meets genetic drift: the coevolution of male traits and female preferences in finite populations

**DOI:** 10.1101/2024.11.16.623954

**Authors:** Kuangyi Xu

## Abstract

Fisher’s mechanism is central to sexual selection theories, where mate choice generates a genetic correlation between male trait and female preference alleles, driving the coevolution of the trait and preference with positive feedback. However, how Fisher’s mechanism operates in finite populations remains unclear, as sexual selection can interact with genetic drift, influencing both trait-preference correlation and allele frequencies. By using population genetic models, this study addresses the gap in our understanding of interactions between fundamental evolutionary forces. We show that more frequent recombination increases trait-preference correlations in infinitely large populations, unless a positive correlation initially exists. In finite populations, interactions between sexual selection and drift elevate trait-preference correlation when the male trait is rare but reduce the correlation when the trait is common, potentially making it negative when recombination is rare or population size is small. Also, these interactions tend to slow the spread of the male trait while promoting the evolution of preferences. These results differ from the Hill-Robertson effect under natural selection due to two key factors: mate choice generates positive linkage disequilibrium, and the strength of indirect selection on preferences increases with linkage disequilibrium. The fixation of trait and preference alleles is positively correlated. This correlation peaks at intermediate recombination rates and is often stronger in small populations than in large ones, so large population sizes tend to reduce the likelihood that both trait and preference alleles fix. We discuss how the results provide insights into the progression of sexual selection in nature.

## Introduction

The evolution of exaggerated display traits in males and why females prefer these seemingly non-adaptive traits have received long-standing attention in evolutionary biology. Fisher (1958) elucidated that males with elaborate display traits may enjoy higher mating success from female preferences, which allows the spread of alleles for elaborate characters. On the other hand, females that prefer to mate with males bearing elaborate traits can be indirectly favored since they tend to produce offspring carrying the alleles for both the attractive display traits and the preference. This process will thus generate positive genetic correlation between male trait alleles and preference alleles and may result in their coevolution towards extreme values, known as Fisherian runaway (Lande 1981; Kirkpatrick 1982; Hall et al. 2000; Kuijper et al. 2012; Fry 2022; Xu et al. 2023). Since then, Fisher’s mechanism has been central to most sexual selection models (Mead & Arnold 2004; Prum 2010).

Despite being theoretically plausible, it remains doubtful whether runaway selection is a powerful force in producing exaggerated traits and strong preferences in nature. Some models and meta-analyses suggest that genetic correlations between male traits and preferences tend to be weak, so that indirect selection on female preferences via genetic correlation is likely to be weaker than direct selection that arises from direct benefits or search costs (Kirkpatrick & Barton 1997; Greenfield et al. 2014; Servedio 2024; but see Fry 2022). Moreover, individual-based simulations suggest that the trait-preference correlation may be unlikely to persist in finite populations of biologically realistic sizes and tends to be overwhelmed by genetic drift (Nichols & Butlin 1989; Roff & Fairbairn 2014; Xu et al. 2023).

To understand the effectiveness of Fisher’s mechanism in nature, a fundamental unresolved question is how sexual selection through mate choice operates in finite populations, as previous sexual selection models often assume infinitely large populations and focus on the equilibrium state. Runaway selection critically depends on the genetic correlation between male traits and female preferences, but this correlation can be influenced by interactions between runaway selection and genetic drift. It is well-established that the interaction between natural selection and drift will generate negative linkage disequilibrium (LD) between two beneficial alleles, causing the alleles to mutually inhibit each other’s sweep—a phenomenon known as the Hill-Robertson effect (Hill & Robertson 1966; Kouyos et al. 2007; Comeron et al. 2008). Similarly, interactions between runaway selection and drift are expected to affect both the sign and magnitude of LD between trait and preference alleles, as well as their allele frequencies.

The mechanism underlying the Hill-Robertson effect has been well studied (e.g., Barton & Otto 2005). Briefly, random sampling in finite populations will generate either positive or negative LD with (nearly) equal probabilities. However, due to larger genetic variation, selection is more efficient when beneficial alleles are in the same haplotype (i.e., positive LD) than in opposite haplotypes (negative LD). Therefore, positive deviations in LD will decay faster than negative deviations. In addition, there will be positive covariance between deviations in allele frequencies and deviations in LD, making positive deviations in LD decay even faster. Consequently, LD will be negative on average and beneficial alleles will inhibit the sweep of each other.

Nevertheless, what we have learned from the Hill-Robertson effect may not directly apply to the case of sexual selection, since sexual selection through mate choice differs from natural selection in several aspects. Importantly, mate choice can generate positive LD between male traits and preferences, which appears to be similar to the generation of positive LD under positive epistasis between loci. However, under sexual selection, more frequent recombination can promote the buildup of LD (Figure 1), while under epistasis, recombination always reduces the level of LD (Kouyos et al. 2007). Moreover, under sexual selection, the selection strength on male traits depends on the frequency of preference alleles, while selection on preference alleles is indirect, with the selection strength being proportional to the selection strength on the male trait and the strength of LD (Lande 1981; Kirkpatrick 1982).

**Figure 1.**
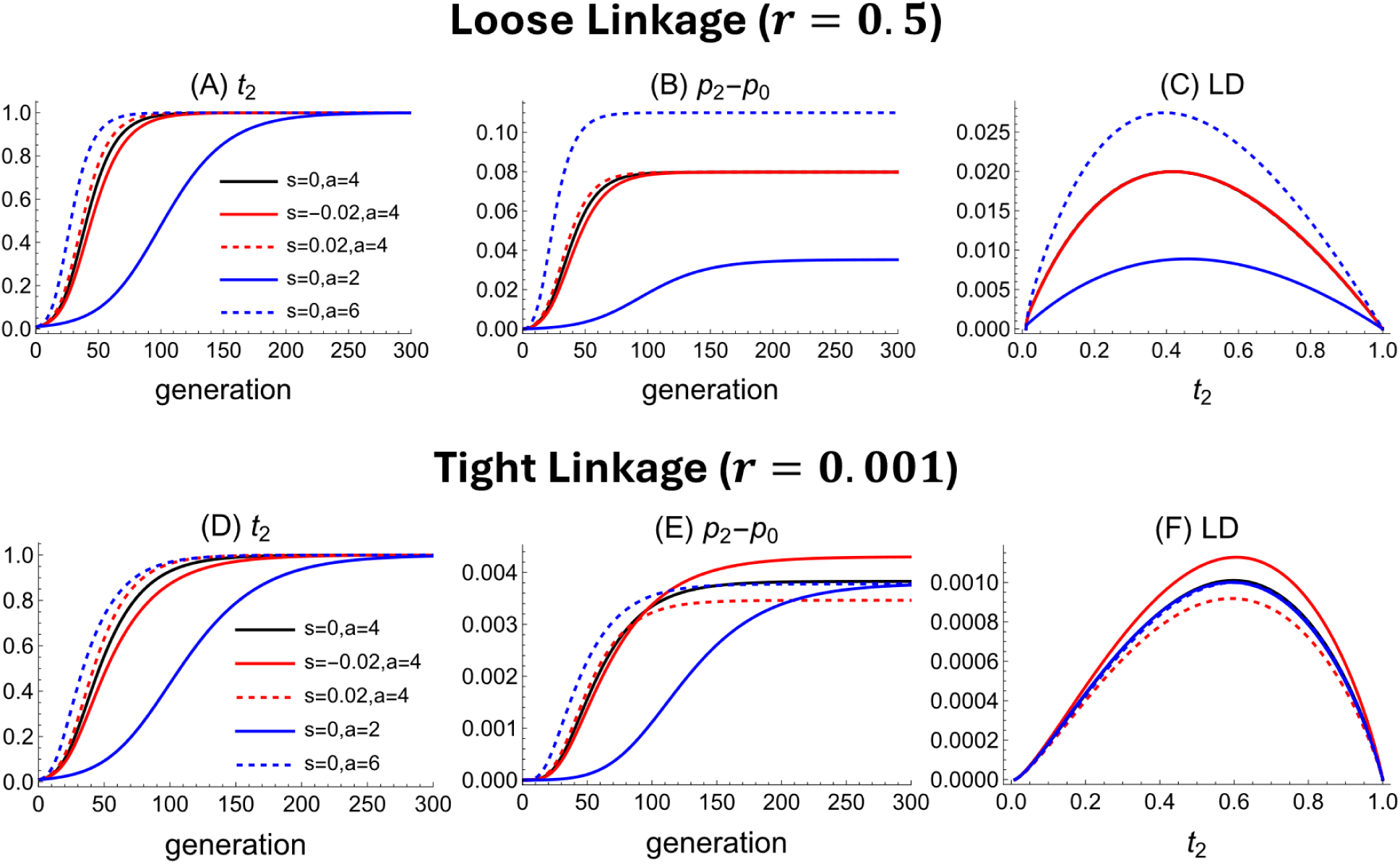
Dynamics of the male trait frequency *t*_2_, preference allele frequency *p*_2_, and linkage disequilibrium (LD) in infinitely large populations. Upper and lower panels show results under loose and tight linkage between male trait and preference loci, respectively. In panels (C) and (F), the results are plotted against the male trait frequency rather than generations to account for different rates of evolution, and the black line overlaps with other lines and thus cannot be seen. Unless otherwise specified, parameters are *s* = 0, *a* = 4, *t*_0_ = 0.01, *p*_0_ = 0.1, *D*_0_ = 0.

To address the gap in our understanding of the interaction between fundamental evolutionary forces, this study investigates how the interaction between sexual selection through mate choice and genetic drift affects the progression of the runaway process in finite populations. Since previous sexual selection models focus on the equilibrium state, we first examine the dynamics of the frequencies of male trait and preference alleles, as well as the LD between them, in infinitely large populations. We find that more frequent recombination increases the magnitude of LD between trait and preference alleles, except when the LD is initially positive.

The interaction between sexual selection and drift increases the trait-preference correlation when the male trait is rare but reduces it when the trait becomes common. This interaction also reduces the frequency of male trait alleles while elevating the frequency of preference alleles. These results contrast with previous findings for natural selection. These differences arise primarily due to two key factors: (1) sexual selection through mate choice generates positive LD, and (2) the strength of indirect selection on preferences increases with LD. Additionally, we examine the effects of key factors (e.g., recombination rate) on the fixation of male trait and preference alleles, which can be understood in light of the above findings.

## Model

We consider a haploid polygynous population with equal sex ratio and a constant population size 2*N*. Following Kirkpatrick (1982), we consider a locus T that governs a trait expressed only in males and a locus P that determines the female mating preference. The T locus has two alleles *T*_1_ and *T*_2_. *T*_1_ males are without the secondary sexual characteristic, while *T*_2_males bear a trait which is more attractive to *P*_2_ females than *T*_1_ males. *P*_1_ females mate indiscriminately, while *P*_2_females prefer to mate with a *T*_2_male *a* times more frequently than with a *T*_1_male. The male trait may be subject to viability selection, and the fitness of *T*_1_and *T*_2_males is 1 and 1 + *s*, respectively. The selection coefficient *s* may be either positive or negative. To focus solely on the sexual selection, we assume no direct viability or fertility selection at the P locus, so there is no Hill-Robertson effect caused by natural selection at the two loci, which could otherwise be confounding.

We census immediately after juveniles have been randomly sampled from the offspring produced by the previous generation. We assume that *N* female and *N* male juveniles are sampled, in order to eliminate fluctuations in effective population size and selection strength on sex-limited traits due to changes in sex ratios caused by random sampling. After the census, the population undergoes viability selection in males, non-random mating via mate choice and offspring reproduction, where we assume that the number of potential offspring is sufficiently large such that these processes can be treated deterministically.

We denote the frequency of *T*_*i*_ and *P*_*i*_ (*i* = 1,2) in the juvenile population before selection by *t*_*i*_ and *p*_*i*_, respectively. We describe the system in terms of allele frequencies of *t*_2_and *p*_2_, and the linkage disequilibrium *D* = *X*_*T*1*P*1_ *X*_*T*2*P*2_ − *X*_*T*1*P*2_ *X*_*T*2*P*1_, where *X*_*i*_ is the frequency of haplotype *i*. The deterministic recursions of *t*_2_, *p*_2_ and *D* in infinitely large populations are given by equation (1) in Kirkpatrick (1982). In finite populations, *t*_2_, *p*_2_and *D* are random variables, and our focus is their average values across replicate populations (i.e., expectation). Throughout the paper, we use 𝔼[*X*] to denote expectation of a random variable *X*.

### Deviations from the deterministic trajectory

In finite populations, random sampling at each generation causes *t*_2_, *p*_2_ and *D* to differ from the values predicted by deterministic recursions in infinitely large populations, and we denote the perturbations that arise from a single generation of sampling by ζ_*t*_ and ζ_*p*_ and ζ_*D*_. The expectation of perturbations in allele frequencies are 𝔼[ζ_*t*_] = 𝔼[ζ_*p*_] = 0, while the expectation of the perturbation in LD is 𝔼[ζ_*D*_] = − D/2N. The magnitude of the second moments of these 2*N* perturbations is at the order of 1/*N* (see equation (A1) in Barton & Otto (2005)). Due to perturbations at each generation, the actual trajectory of *t*_2_, *p*_2_and LD in a finite population will deviate from the deterministic trajectory, and we denote the deviations in *t*_2_, *p*_2_ and LD by δ*t*, δ*p* and δ*D*, respectively.

We follow the perturbation analysis in Barton & Otto (2005) to obtain the expectation of deviations. The analysis assumes that the deviations remain small, and the population size is large enough so that we can focus on the first and second moments of the deviations, which are of order *N*^−1^, and ignore *O*(*N*^−2^) terms. We first obtain the recursions for the three deviations based on the deterministic recursions of allele frequencies and LD given in Kirkpatrick (1982). Briefly, let *x*^∗^ = *f*(*x*) be the allele frequencies and LD after one generation based on the deterministic recursions, where *x* = (*t*_2_, *p*_2_, *D*). The recursion of the deviation vector *z* = (δ*t*, δ*p*, δ*D*) is given by *z*^∗^ = *f*(*x* + *z*) − *f*(*x*) + ζ, where the vector ζ = (ζ_*t*_, ζ_*p*_, ζ_*D*_) represents the perturbations described above. Based on the recursion of deviations, we can obtain the recursions of the first moments of the deviations (see Appendix), given by

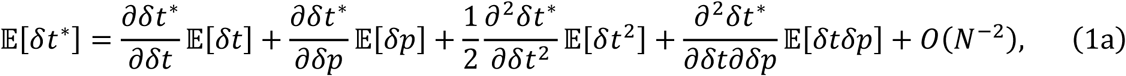

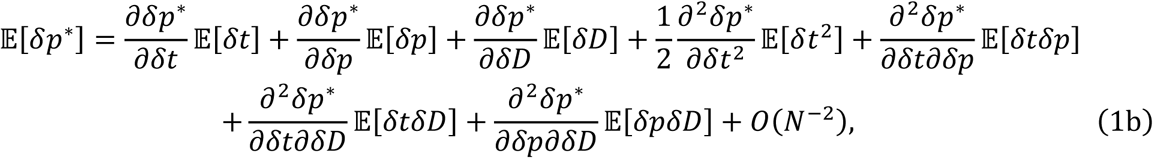

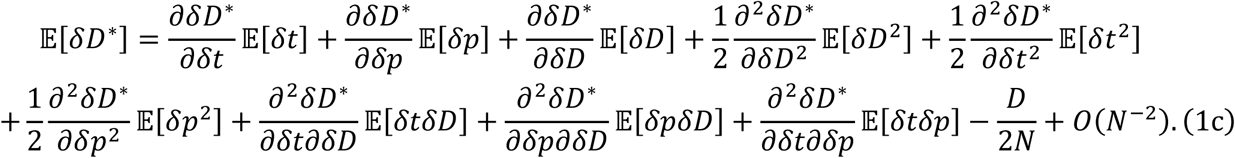

The second moments of deviations include the variance in the three deviations, 𝔼[δ*t*^2^], 𝔼[δ*p*^2^] and 𝔼[δ*D*^2^], as well as the covariance between deviations 𝔼[δ*t*δ*p*], 𝔼[δ*t*δ*D*] and 𝔼[δ*p*δ*D*]. The recursions of the second moments are presented in the Appendix. Detailed expression for the recursions of deviations as well as their first and second moments are available in the Mathematica Notebook available at https://doi.org/10.5281/zenodo.14172252.

Unfortunately, since the recursions of deviations in LD and allele frequencies are mutually dependent (see Equations (1) and (S4)), making it impossible to apply the quasi-linkage equilibrium approximation (Kimura 1965; Nagylaki 1993), which assumes that terms involving deviations in LD reach a steady state on a faster timescale than deviations in allele frequencies.

Therefore, we must numerically solve all nine recursions involving the first and second moments of the deviations. The predictions from the perturbation analyses are then compared to the averages across replicates obtained from simulations.

## Results

We consider the scenario when the preference alleles are segregating, and the male trait allele *T*_2_will sweep to fixation in infinitely large populations through sexual selection or/and natural selection. We first examine the dynamics of allele frequencies and LD in infinitely large populations. We then show how the interaction between sexual selection and genetic drift in finite populations generates deviations from the deterministic dynamics and investigate the underlying mechanisms. Finally, we examine the fixation probability of trait and preference alleles as well as the correlation in fixation events.

### Deterministic dynamics of male trait and preference

Kirkpatrick (1982) clarifies that the selection strength on the male trait allele *T*_2_depends on the frequency of *P*_2_ females, while the selection strength on *P*_2_is proportional to both LD and the selection strength on *T*_2_. When the benefit from sexual selection outweighs the (potential) viability cost associated with the male trait, the male trait allele *T*_2_ will sweep to fixation, which also leads to an increase in the preference allele frequency through positive LD generated by mate choice (see Figure 1).

The impacts of recombination rate on LD between trait and preference alleles depends on the initial LD. When trait and preference alleles are initially at linkage equilibrium, a higher recombination rate between them can significantly increase the magnitude of linkage disequilibrium (LD) (compare the magnitude of LD in Figures 1C and 1F), which in turn amplifies the increase in the preference allele frequency (compare Figures 1B and 1E). This is because recombination facilitates the formation of haplotype *T*_2_*P*_2_, and once *T*_2_*P*_2_is formed, it is unlikely to be decoupled by recombination under mate choice, as *T*_2_*P*_2_females mate disproportionally often with males carrying allele *T*_2_, and *T*_2_*P*_2_males are more likely to be chosen by *P*_2_females. In other words, the benefit of increased recombination in forming *T*_2_*P*_2_outweighs the cost of destroying *T*_2_*P*_2_under mate choice. However, if LD between trait and preference alleles is initially positive, tighter linkage will increase the magnitude of LD, resulting in a larger increase in the preference allele frequency (Figure S1).

Although stronger selection on the male trait can result from either higher viability or stronger preferences (Figures 1A, 1D), we may distinguish between the two causes by examining the magnitude of LD and changes in the preference allele frequency. Specifically, male trait viability is more effective in affecting the magnitude of LD and the overall increase in the preference allele frequency when the linkage between the male trait and preference loci is tighter (compare *s* = −0.02, 0, 0.02 in Figures 1C vs. 1F and 1B vs. 1E). In contrast, the impact of preference strength is more significant when linkage is looser (compare *a*=2, 4, 6 in Figures 1C vs. 1F, 1B vs. 1E).

The differences above arise because viability and sexual selection on the male trait influence LD in different ways. Specifically, reduced viability of the male trait *T*_2_ increases the magnitude of LD by slowing the sweep of *T*_2_, as a slower sweep allows LD to accumulate over more generations. Therefore, when recombination is more frequent, the impact of the sweep rate—and thus the viability of the male trait—on LD becomes weaker, since LD will approach equilibrium more rapidly. In contrast, stronger preference strength directly increases LD by facilitating the generation of haplotype *T*_2_*P*_2_through recombination, as explained before. Therefore, stronger preference strength is more effective at elevating LD when recombination is more frequent.

The coevolution of trait and preference is also influenced by the initial frequencies of the male trait and female preference alleles. A lower initial frequency of the male trait *T*_2_, by causing a slower sweep of *T*_2_, elevates the magnitude of LD and the increase in the preference allele frequency (Figure S1), and this impact is more significant when linkage is tighter. The initial frequency of the preference allele *P*_2_, like preference strength, influences not only the rate of sweep but also the generation of LD. Therefore, a higher initial frequency of *P*_2_may reduce the magnitude of LD and the increase in *P*_2_ frequency under tight linkage due to a faster sweep of the male trait, but tends to elevate both quantities when linkage is loose (Figure S2).

### Deviation of allele frequencies and LD in finite populations

In finite populations, deviations in both allele frequencies and LD change non-monotonically over time. Specifically, as the male trait allele *T*_2_ sweeps from low to high frequency, the expected deviation in LD, 𝔼[δ*D*], is initially positive, then declines to be negative, increases again and eventually converges toward 0 (Figure 2A). The dynamics of the expected deviation in the frequency of the male trait, 𝔼[δ*t*], follow a similar pattern (Figure 2B). Unlike 𝔼[δ*D*] and 𝔼[δ*t*], the expected deviation in the frequency of the preference allele, 𝔼[δ*p*], does not converge to 0, and will remain positive when linkage is sufficiently tight (Figure 2C).

**Figure 2.**
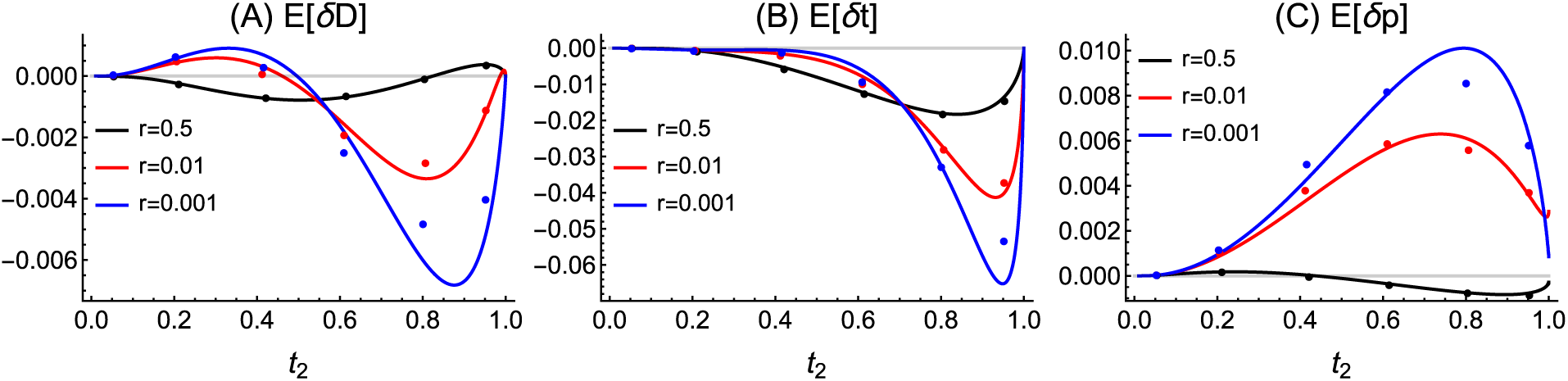
Average deviations from the deterministic trajectory of *t*_2_, *p*_2_ and LD in finite populations under different recombination rates. Deviations are plotted against *T*_2_frequency from the deterministic trajectory (rather than against generation) to account for different rates of evolution under different recombination rates. Lines are predictions calculated from Equation (1), and dots are the average values calculated from 10^5^simulation replicates. Parameters used are 2*N* = 10000, *a* = 4, *t*_0_ = 0.01, *p*_0_ = 0.1, *D*_0_ = 0. For *r* = 0.001, *T*_2_ will be lost in some simulation replicates, so the simulation results deviate from the model prediction.

In general, the magnitude of deviations in allele frequencies and LD becomes larger when recombination rate is lower (compare lines with different *r* values in Figure 2), since perturbations from earlier generations decay more slowly, thereby accumulating to create larger deviations in later generations. Consequently, under loose linkage, the expected dynamics of allele frequencies and LD in finite populations will closely resemble the dynamics observed in infinitely large populations (compare the magnitude of deviations in Figure 2 and the dynamics in Figure 1 at *r*=0.5), unless the population size is sufficiently small (Figure S3). In contrast, under tight linkage, the sweep of *T*_2_is significantly slowed (*r*=0.001 in Figure 2B). Also, the expected dynamics of LD and *P*_2_ frequency will be nearly identical to the dynamics of their deviations, since the magnitude of deviations is much larger than the changes observed under the deterministic case (compare results at *r*=0.001 between Figures 2A vs1C, and Figures 2C vs 1B). The magnitude of deviations also become larger when the male trait allele sweeps more slowly (Figure S4) and the initial LD is higher (Figure S5).

The perturbation analysis assumes that the population size is large enough so that *T*_2_ will reach fixation. However, when the population size is small so that *T*_2_ may be lost due to drift, the expected deviations in allele frequencies and LD will depend on whether *T*_2_ becomes fixed or lost (Figure S3). In replicates where *T*_2_becomes fixed, the deviation in LD is initially positive, with a magnitude much larger than that observed in large populations. In contrast, in replicates where *T*_2_is lost, the average deviation in LD remains negative over time.

### Biological explanations

We investigate the mechanisms underlying the complicated dynamics of deviations in LD and allele frequencies, by calculating the contribution of components in Equation (1). We focus on the dominant components that make major contribution to these deviations under two extreme scenarios: when linkage is very loose and when it is very tight. The results for an intermediate recombination rate fall between those observed under the two extremes.

### Deviation in LD under loose linkage

Under loose linkage, the deviation in LD, 𝔼[δ*D*], is primarily driven by three components (bolded lines in Figure 3A): 1) the deviation in the frequency of male trait *T*_2_, 𝔼[δ*t*], 2) the variance of the deviation in *T*_2_ frequency, 𝔼[δ*t*^2^], and 3) the covariance between deviations in the frequencies of *T*_2_ and *P*_2_, 𝔼[δ*t*δ*p*].

**Figure 3.**
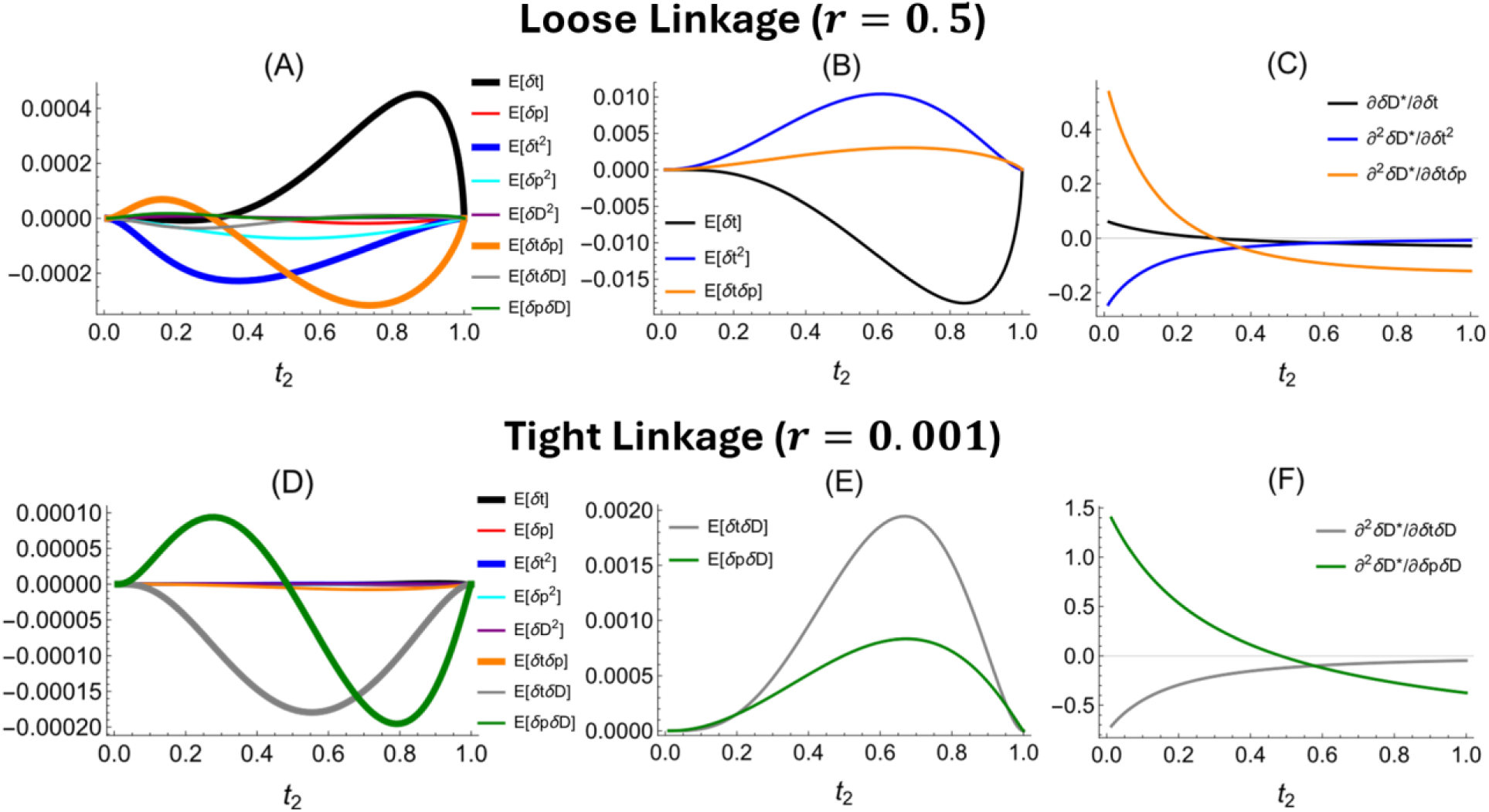
Analysis of the mechanism underlying the deviation in LD, 𝔼[δ*D*], for loose linkage (upper panels) and tight linkage (bottom panels). First column: contributions from each component in Equation (1.3) to 𝔼[δ*D*]. Curves are bolded for dominant terms. Second column: dynamics of the values of the dominant terms. Third column: effects of the dominant terms on 𝔼[δ*D*]. Other parameters used are the same as those in Figure 2.

Since LD initially builds up and then decline as the male trait alleles sweeps (Figure 1C), the deviation in *T*_2_frequency, 𝔼[δ*t*], contributes positively to the deviation in LD when *T*_2_ is rare (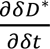 > 0), but negatively when *T*_2_ becomes common (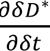 < 0). However, 𝔼[δ*t*] changes from positive when *T*_2_is rare to negative as *T*_2_becomes common (Figure 3B; the reasons are explained later). Therefore, 𝔼[δ*t*] tends to increase the deviation in LD throughout times (black line in Figure 3A).

The variance 𝔼[δ*t*^2^] always reduces the deviation in LD (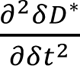 < 0; Figure 3C). This is because LD is a concave function of the male trait frequency *t*_2_(Figure 1C). Therefore, when *T*_2_ is rare, so that an increase in its frequency *t*_2_promotes the buildup of LD, the increase in LD caused by a positive deviation in *t*_2_is smaller than the decrease in LD caused by a negative deviation in *t*_2_. When *T*_2_ is common, in which case an increase in *T*_2_frequency accelerates the decay of LD, the additional decay in LD resulting from a positive deviation in *t*_2_outweighs the slower decay under a negative deviation in *t*_2_.

Since positive feedback in the coevolution of trait and preference alleles, the covariance between deviations in allele frequencies, 𝔼[δ*t*δ*p*], remains positive (Figure 3B). Similar to 𝔼[δ*t*], the effect of 𝔼[δ*t*δ*p*] on the deviation in LD changes from positive to negative as the frequency of *T*_2_increases (Figure 3C). This means that the increase in LD resulting from positive deviations in both *T*_2_ and *P*_2_ frequencies is larger than the reduction in LD caused by negative deviations in the frequencies of both alleles (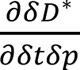 > 0). When *T*_2_ becomes common, the additional decay in LD from positive deviations in *T*_2_ and *P*_2_frequencies outweighs the slowing of LD decay when both deviations are negative.

### Deviation in LD under tight linkage

Unlike the results under loose linkage, when linkage is tight, terms involving only deviations in allele frequencies (e.g., 𝔼[δ*t*], 𝔼[δ*t*δ*p*]) are no longer the primary contributors to the deviation in LD. Instead, the deviation in LD is mainly driven by the covariances between deviations in LD and allele frequency, 𝔼[δ*t*δ*D*] and 𝔼[δ*p*δ*D*] (bolded lines in Figure 3D). This is because deviations in LD decay more slowly than deviations in allele frequencies when linkage is tight. However, the contribution from the variance of deviations in LD, 𝔼[δ*D*^2^], remains small (Figure 3D).

The covariance between deviations in *T*_2_frequency and LD, 𝔼[δ*t*δ*D*], stays positive throughout the sweep (Figure 3E). This covariance always reduces the deviation in LD (i.e., 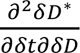 < 0; Figure 3F), similar to the result found for the Hill-Robertson effect under natural selection (Barton & Otto 2005). Biologically, in populations that happen to have positive deviations in both LD and *T*_2_frequency (δ*D*, δ*p* > 0), this positive δ*D* will decay faster in future generations than the decay of a negative δ*D* in populations that happen to have negative δ*D* and δ*p*.

Similarly, the covariance between deviations in *P*_2_ frequency and LD, 𝔼[δ*p*δ*D*], remains positive (Figure 3E). This is because selection strength on *P*_2_ is proportional to the magnitude LD, so a positive deviation in LD will accelerate the evolution of *P*_2_, resulting in a positive deviation in *P*_2_ frequency. However, the effect of this covariance on the deviation in LD, 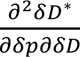, declines from positive to negative as *T*_2_ increases in frequency (Figure 3F).

### Deviation in trait and preference allele frequencies

As Equation (1) shows, the deviation in the male trait frequency, 𝔼[δ*t*], is contributed by three factors: 1) the deviation in the frequency of preference allele *P*_2_, 𝔼[δ*p*], 2) the variance of deviations in *T*_2_ frequency (𝔼[δ*t*^2^]), and 3) the covariance between deviations in the frequency of *T*_2_and *P*_2_, 𝔼[δ*t*δ*p*]. The component 𝔼[δ*p*] becomes an important force only under tight linkage, while the other two factors are always dominant forces.

When linkage is loose, the deviation in preference allele frequency, 𝔼[δ*p*], remains nearly 0 (*r*=0.5 in Figure 2C), contributing minimally to the deviation in male trait frequency 𝔼[δ*t*] (Figure 4A). In contrast, when linkage is tight, 𝔼[δ*p*] becomes large and stays positive (*r*=0.001, Figure 2C), contributing positively to 𝔼[δ*t*] (Figure 4C) since a higher *P*_2_ frequency accelerates the evolution of *T*_2_ by increasing the selective advantage (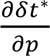 > 0).

**Figure 4.**
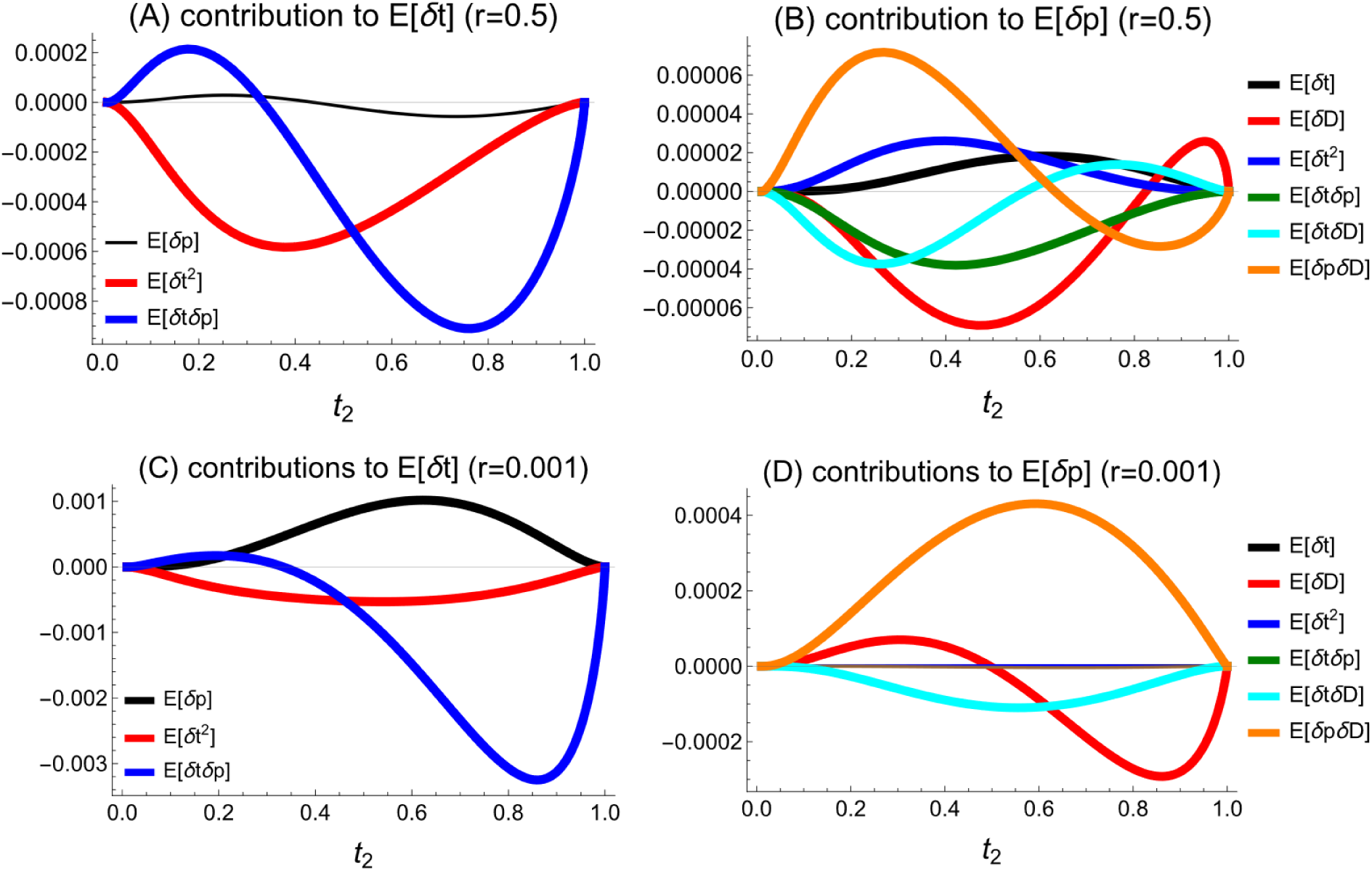
Contributions of components to the deviation in male trait frequency 𝔼[δ*t*] and female preference frequency 𝔼[δ*p*], calculated from Equations (1.1) and (1.2), respectively. Upper and bottom panels show results under loose and tight linkage, respectively. Other parameters used are the same as those in Figure 2.

The variance of deviations in *T*_2_ frequency, 𝔼[δ*t*^2^], reduces the deviation in *T*_2_ frequency (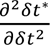 < 0; Figures 4A, 4C). This means that a positive deviation in *T*_2_ frequency decays faster in future generations than a negative deviation in *T*_2_ frequency.

The covariance between deviations in allele frequencies, 𝔼[δ*t*δ*p*], remains positive, as explained before. However, its effect on 𝔼[δ*t*] declines from positive to negative as the frequency of the male trait *T*_2_ increases (Figures 4A, 4C). Intuitively, positive deviations in both *T*_2_and *P*_2_ frequencies are more efficient at driving the increase of *T*_2_ frequency when *T*_2_ is rare than when *T*_2_ is already common.

The deviation in preference allele frequency is mainly contributed by terms involving deviations in LD, including 𝔼[δ*D*], 𝔼[δ*p*δ*D*] and 𝔼[δ*t*δ*D*] (Figures 4B, 4D), since selection on preference occurs indirectly through LD. However, when linkage is very loose, terms involving the deviation in male trait frequency, δ*t*, can also play some roles in contributing to 𝔼[δ*p*] (Figure 4B), because the selection strength on *P*_2_ is proportional to the selection strength on the male trait.

### Fixation of male trait and preference alleles

This section investigates the impacts of genetic and ecological factors on the probability of fixation of the male trait allele *T*_2_ and the female preference allele *P*_2_. To understand the likelihood of runaway, we also assess the probability that both *T*_2_ and *P*_2_ become fixed. We consider the case in which a single mutant *T*_2_at the male trait locus occurs in a population with alleles *P*_1_and *P*_2_segregating at the preference locus.

### Effects of recombination

In general, the fixation probability of the male trait allele *T*_2_ increases as the recombination rate between trait and preference loci is higher (Figure 5A). This pattern can be understood by referring to Figure 2B, where tighter linkage tends to slow the sweep of *T*_2_ in finite populations (i.e., more negative 𝔼[δ*t*]).

**Figure 5.**
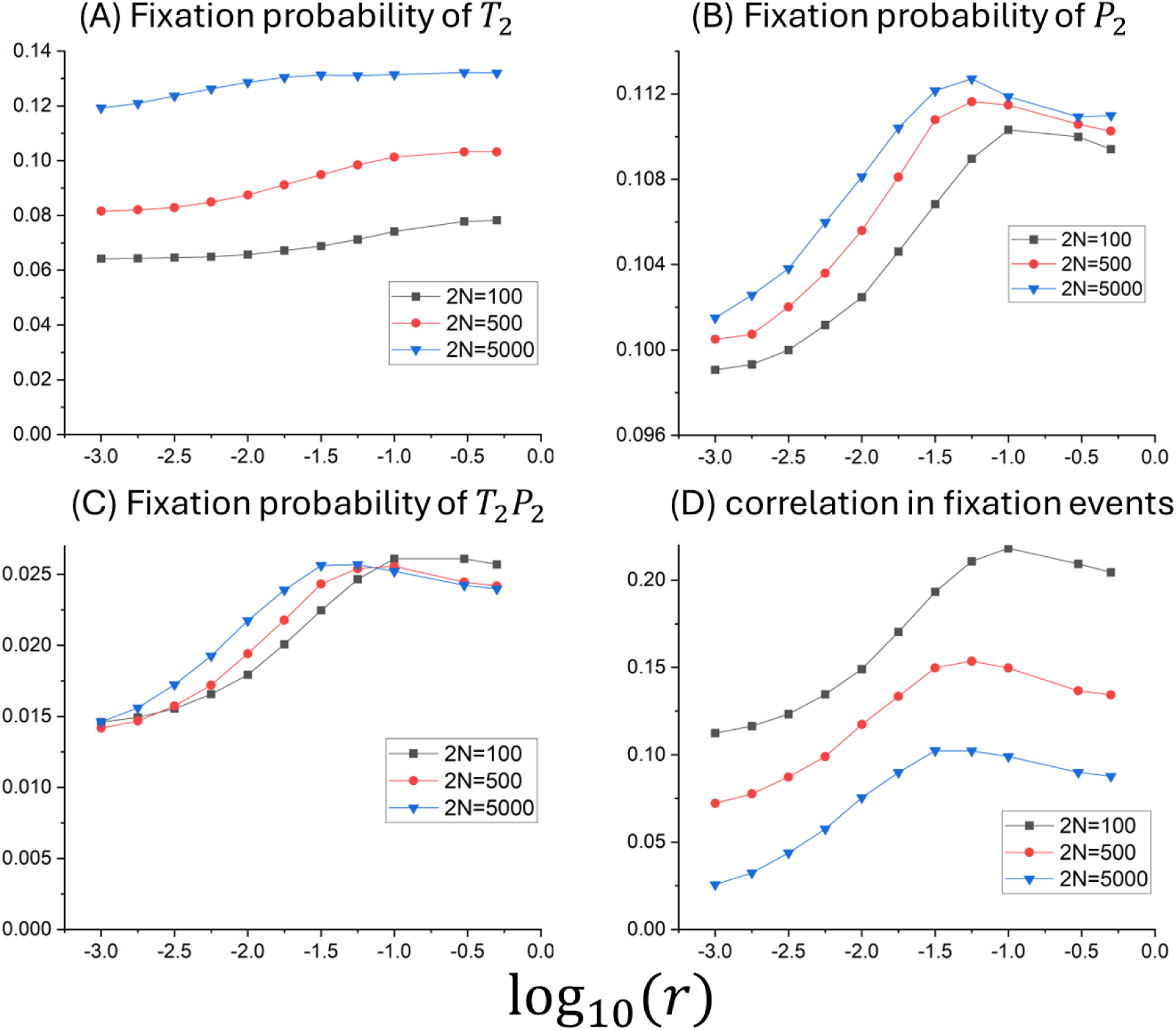
Effects of recombination rate *r* (x axis) on the fixation probability of a single mutant *T*_2_at the male trait locus and the fixation probability of allele *P*_2_at the preference locus. The initial frequency of *P*_2_ is fixed to be *p*_0_ = 0.1. Panel (D) shows the correlation between the fixation probability of *T*_2_and *P*_2_. Other parameters are *a* = 4, *s* = 0.

In contrast, the fixation probability of the preference allele *P*_2_ peaks at an intermediate recombination rate, with this optimal rate shifting higher in smaller populations (Figure 5B). To understand this, recall that the expected change of *P*_2_frequency in finite populations can be decomposed into the deterministic change in infinitely large populations and the deviation 𝔼[δ*p*]. Under loose linkage, the deterministic increase is substantial, but the deviation is nearly zero (compare results at *r* = 0.5 in Figures 1B and 2C). Conversely, under tight linkage, the deterministic increase is small, while the deviation is large and tends to be positive (Figure 2C). Therefore, an intermediate recombination rate balances the two components, maximizing the expected increase of *P*_2_frequency.

The fixation of trait and preference alleles is always positively correlated. This correlation in fixation events, and consequently the probability that both trait and preference alleles fix, peaks at an intermediate recombination rate (Figures 5C, 5D). For intuition, the correlation in fixation events is primarily determined by the initial LD between trait and preference alleles during their coevolution. A higher recombination rate increases the deterministic generation of initial LD (Figure 1), while a lower recombination rate elevates initial LD through the interaction between sexual selection and drift (Figure 2A). Therefore, an intermediate recombination rate maximizes the initial LD by balancing the two contributions.

### Effects of population size

The fixation probability of a single mutant allele *T*_2_ is lowest at an intermediate population size (Figure 6A). Intuitively, a smaller population size increases the reduction in *T*_2_ frequency caused by the interaction between sexual selection and drift, since the deviation in *T*_2_frequency, 𝔼[δ*t*], is proportional to 1/*N*. However, when the population size is excessively small, drift dominates over selection in determining the fate of *T*_2_, and the fixation probability approaches that of a neutral mutant (i.e., 1/4*N*), which increases in smaller populations. Therefore, the population size that minimizes the fixation probability of *T*_2_ shifts lower when the male trait is more strongly favored (Figures 6A, S6 and S7).

**Figure 6.**
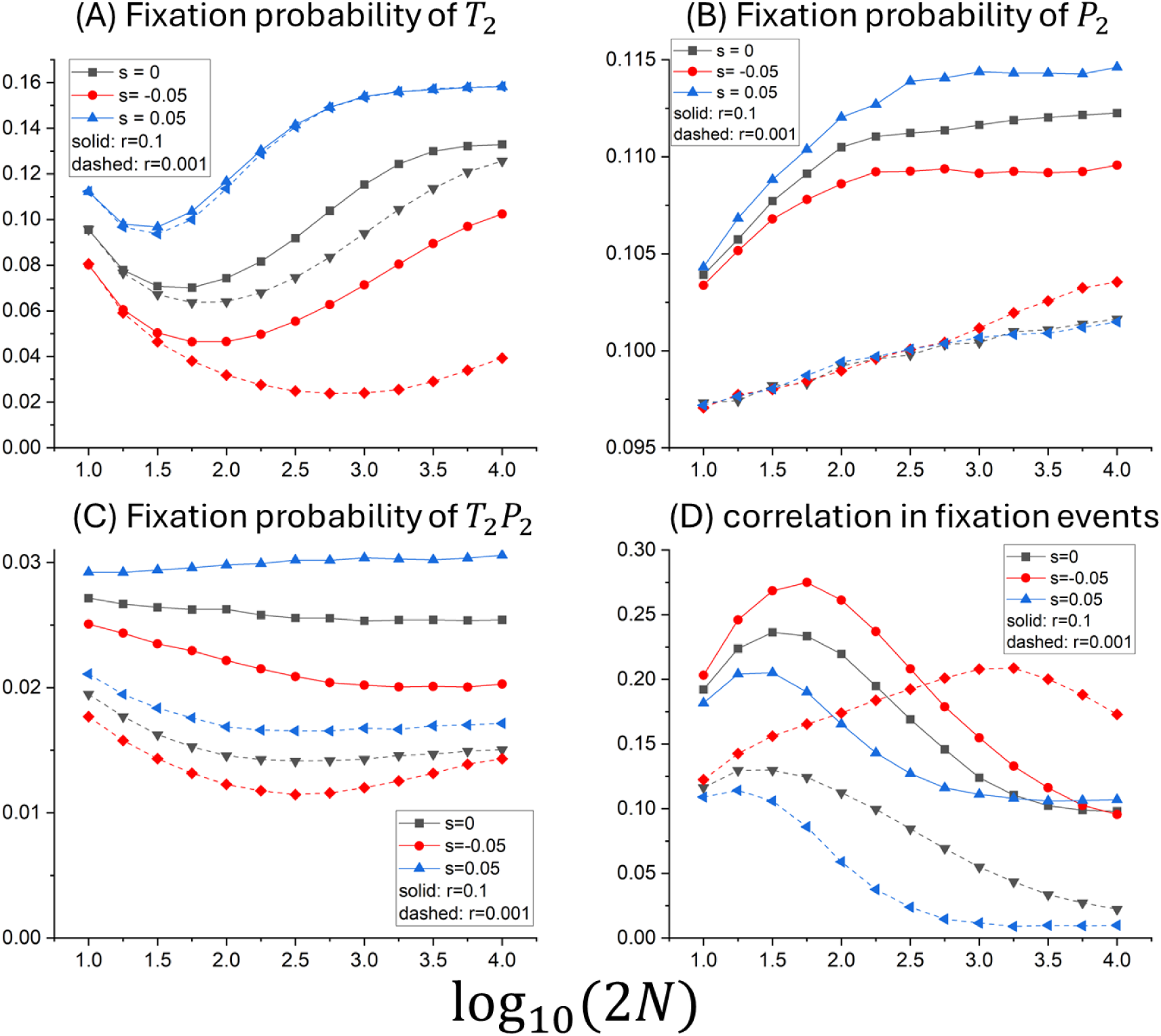
Effects of population size (x axis) and selection coefficient on the male trait on the fixation probability of a single mutant *T*_2_at the male trait locus and the fixation probability of allele *P*_2_at the preference locus. Panel (D) shows the correlation between the fixation probability of *T*_2_and *P*_2_. Other parameters are *p*_0_ = 0.1, *a* = 4, *s* = 0.

The fixation probability of preference allele *P*_2_ is generally higher in larger populations (Figure 6B), since we assume a fixed initial frequency of *P*_2_. Interestingly, under tight linkage between trait and preference alleles, the fixation probability of *P*_2_can fall below its initial frequency *p*_0_ in small populations (see dashed lines at log_10_(2*N*) < 2.5 in Figure 6B). This is because the trait-preference genetic correlation tends to be negative when recombination is infrequent and drift is strong (Figure S3).

While the fixation probability of both trait and preference alleles is higher in large populations compared to small ones (Figure 6A-B), the probability that both alleles fix is often lower in larger populations (Figure 6C), due to a weaker correlation in the fixation of trait and preference alleles (Figure 6D). This is because large populations weaken the elevation of the initial LD between trait and preference alleles caused by the interaction between sexual selection and drift.

### Effects of selection strength on male trait

Although deterministic models show that the strength of indirect selection on preference increases with the selection strength on the male trait, when recombination is rare, the fixation probability of the preference allele actually increases with a weaker selective advantage of the male trait (compare dashed lines in Figure 6B). This is because a slower sweep of the male trait under weaker selection allows LD to accumulate more (Figure S4), resulting in stronger selection on preference. For the same reason, weaker selection on the male trait also increases the correlation in the fixation of trait and preference alleles, making it stronger (Figure 6D).

### Selection-mutation-drift balance

Previous sections focus on the scenario where the male trait undergoes a selective sweep. Here, we investigate the case where the male trait locus remains polymorphic under the selection-mutation-drift balance. Overall, the results are consistent with those observed under the sweep scenario. We assume equal mutation rates between alleles *T*_1_and *T*_2_. Since indirect selection on preference is weak, to avoid the loss of polymorphism at the preference locus by drift, we fix the frequency of the preference allele *P*_2_to be constant. At selection-mutation balance in infinitely large populations, the frequency of the male trait is independent of the recombination rate. LD between trait and preference alleles is always positive, but reduces as recombination rate becomes lower (Figure 7).

**Figure 7.**
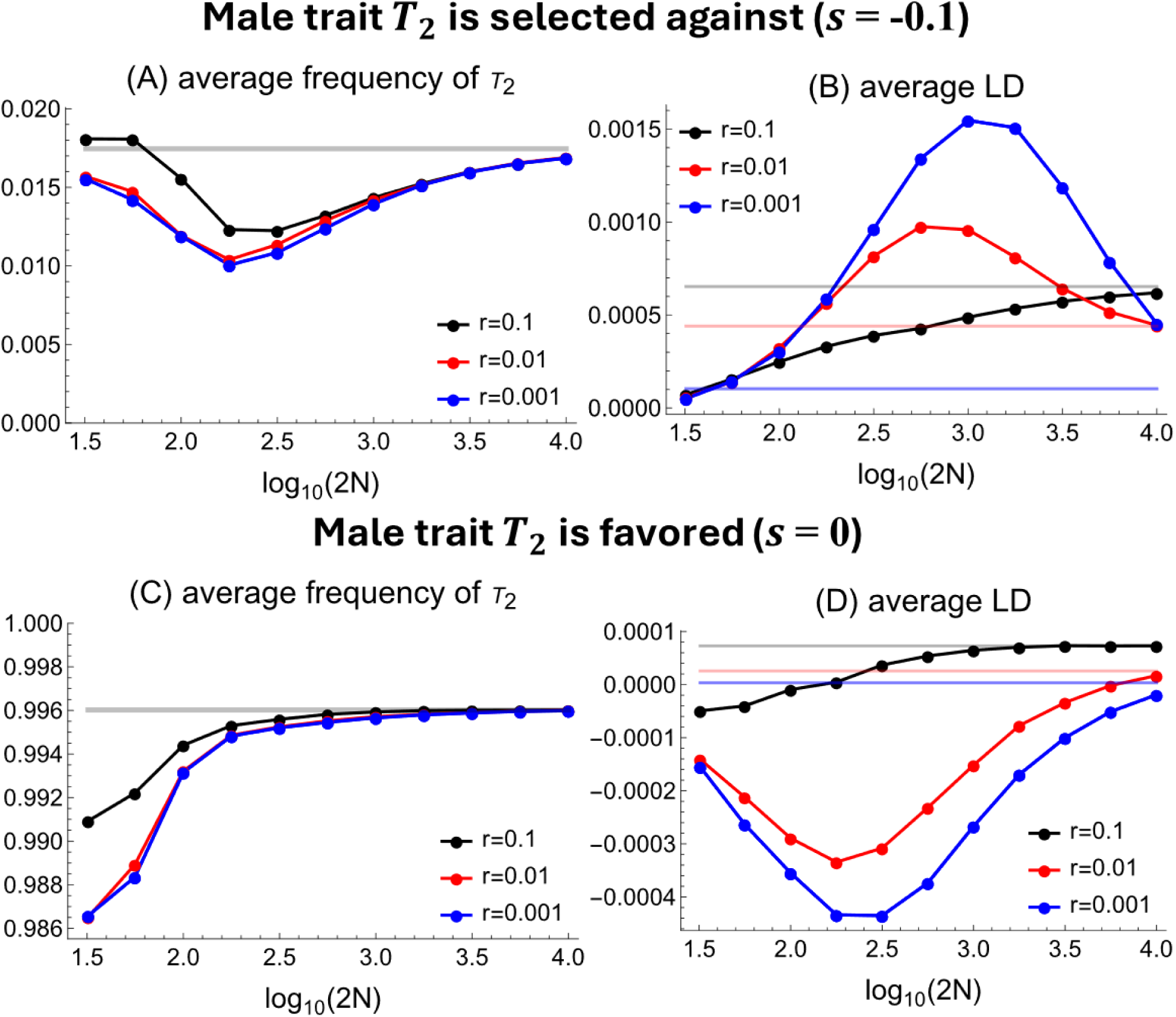
Impacts of population size on the average frequency of the male trait *T*_2_and LD when the male trait locus is at the selection-mutation-drift balance. The allele frequency at the preference locus is fixed at *p*_0_. Horizontal lines denote the equilibrium value in infinitely large populations. Dots are results from 10^6^simulation replicates, where run each replicate for 12*N* generations, starting from the equilibrium state in infinitely large populations. Parameters used are *a* = 2, *p*_0_ = 0.1, μ = 10^−4^.

In finite populations, the average frequency of the male trait at selection-mutation-drift balance tends to be lower than that in infinitely large populations (Figures 7A, 7C), aligning with the finding in Figure 2B. The average male trait frequency decreases with smaller population sizes, except when the population size is extremely small, where drift dominates over selection, causing the trait frequency to approach 0.5 due to the assumption of equal mutation rates between alleles.

Whether the average LD between trait and preference in finite populations is lower or higher than in infinitely large populations depends on the frequency of the male trait allele *T*_2_, consistent with the finding in Figure 2A. Specifically, when *T*_2_is maintained at low frequencies, LD tends to be higher than in infinitely large populations when linkage is sufficiently tight (*r*=0.01, 0.001 in Figure 7B), and LD will be lower than in infinitely large populations when linkage is loose (*r*=0.1 in Figure 7B). When *T*_2_ is maintained at high frequencies, LD is lower than in infinitely large populations and is often negative (Figure 7D). The average LD often reaches its maximum or minimum value at intermediate population sizes, as LD approaches zero in very small populations due to strong genetic drift.

## Discussion

This study investigates the interaction between sexual selection through mate choice and genetic drift during the coevolution of male traits and female preferences in finite populations. Mate choice can directly generate a positive genetic correlation between male trait and preference alleles. The interaction between sexual selection and drift can increase the level of this correlation when male trait alleles are rare, but it tends to reduce the correlation when male trait alleles become common, even making it negative when recombination is infrequent, or population size is small. The interaction between sexual selection and drift also causes deviations in allele frequencies, generally reducing the frequency of male trait alleles while elevating the frequency of preference alleles over the values in infinitely large populations.

The fixation events of male trait and preference alleles are always positively correlated. This correlation tends to be maximized at intermediate population sizes and recombination rates, as the trait-preference genetic correlation at the initial stage of coevolution tends to be low when either the population size or recombination rate is excessively low or high. Therefore, although larger population sizes often increase the fixation probability of both trait and preference alleles, they may reduce the likelihood that both alleles fix.

### Similarity and difference between sexual and natural selection in their interactions with drift

As noted in the introduction, sexual selection through mate choice differs from natural selection in several key aspects. While natural selection can generate LD between loci with epistatic fitness interactions, the interaction between natural selection and drift consistently leads to a negative deviation in LD, regardless of the sign of epistasis (Kouyos et al. 2007). This differs from the results observed under sexual selection, which can be understood by extending the principles underlying the Hill-Robertson effect from natural to sexual selection.

The key principle underlying the observed difference is that the coevolution of male trait and preference alleles proceeds with positive feedback at the initial stage: the selective advantage of preference alleles increases with the selection strength on the male trait and LD, while the buildup of LD also depends on the frequency of trait and preference alleles (Kirkpatrick 1982).

Therefore, positive deviations in LD, by promoting the buildup of LD in later generations, tend to persist longer than negative deviations. However, as male trait alleles become more common, this positive feedback weakens, so positive deviations in LD tend to decay more rapidly than negative deviations, aligning with the mechanisms underlying the Hill-Robertson effect (Barton & Otto 2005). Indeed, when we consider the sweep of two beneficial alleles with positive epistasis, and assume that the selection coefficient of one allele is proportional to the LD between them, we find that deviations in allele frequencies and LD under natural selection behave similarly to those under sexual selection (Figure S8).

The current model focuses on the case when females either mate randomly or exhibit preference. We expect that the results will qualitatively hold in scenarios where females consistently prefer a male trait but with varying strengths (Kirkpatrick 1982), or where females with different preference alleles prefer males with different traits. In both scenarios, Fisher’s mechanism operates similarly, leading to positive feedback during the coevolution of male traits and preferences.

### Detection of trait-preference genetic correlation

Despite the importance of genetic correlation in mediating the coevolution of male traits and preferences (Lande 1981; Kirkpatrick 1982; Mead & Arnold 2004), meta-analyses have found that the level of trait-preference correlation is generally weak (Greenfield et al. 2014). One possible explanation is that the Fisher process may often occur intermittently, with LD disappearing once preference and trait alleles become fixed. The current results provide an additional possible explanation. Specifically, when multiple male trait and preference alleles are either maintained at stable polymorphism or undergoing sweeps, trait-preference correlation may have opposite signs for alleles at different frequencies in finite populations, resulting in an overall weak genetic correlation.

The current results also provide a nuanced understanding of the proposition that lower recombination between trait and preference loci may promote the maintenance of trait-preference correlation and enhance the effectiveness of runaway selection (Otto 1991; Trickett & Butlin 1994; Takimoto et al. 2000). We find that tighter linkage does enhance trait-preference correlation when male trait alleles are maintained at low-frequency polymorphism (Figure 7B), or if trait and preference genes are initially positively linked (Figure S5). The latter condition may occur for trait and preference genes involved in species recognition during speciation (Butlin & Ritchie 1989; Boake 1991), which have been found to co-localize in the same genomic regions in several taxa (Wiley et al. 2012; McNiven & Moehring 2013; Xu & Shaw 2019; Ritchie & Butlin 2024). However, tight linkage can also inhibit the generation of trait-preference correlation when trait and preference genes are initially unlinked or when trait alleles are maintained at high frequency.

### Effects of sexual selection on adaptation

Sexual selection is thought to have mixed effects on adaptation (Candolin & Heuschele 2008; Martínez-Ruiz & Knell 2017). When the male trait is an honest signal of genetic quality (Iwasa et al. 1991), sexual selection can skew reproductive success towards fitter males, thereby increasing the selective advantage of adaptive alleles (Lorch et al. 2003) and helping purge deleterious mutations (Whitlock 2000; Lumley et la. 2015). Our results show that this holds when the population size is large; however, if preference alleles are segregating, sexual selection may inhibit the fixation of adaptive alleles in small populations, particularly when recombination between trait and preference alleles is rare (Figure 6A). Given that sexual selection may reduce the effective population size because only a subset of males may reproduce (Kokko & Brooks 2003), the census population size above which sexual selection will promote the fixation of adaption may be quite large.

### Direct versus indirect selection on preferences

Our models assume no direct selection at the preference locus to eliminate selective interference between trait and preference loci. In reality, female preferences may be subject to direct selection, and whether this direct selection is much stronger than indirect selection due to genetic correlation remains debated (Pomiankowski 1987; Kirkpatrick & Barton 1997; Hall et al. 2000; Kirkpatrick & Hall 2004; Fry 2022; Servedio 2024). However, previous models often assume infinitely large populations. Our results show that the interaction between sexual selection and drift can increase the strength of indirect selection on preferences at intermediate recombination rates, but this effect tends to be quite weak (Figure 5B). Moreover, in finite populations, direct selection at both trait and preference loci can also lead to the Hill-Robertson effect, affecting the genetic correlation and selection strength. If both the preference and the preferred male trait are costly, the Hill-Robertson effect can reduce their correlation. In contrast, if the preference is costly but the male trait is selectively advantageous, the Hill-Robertson effect can increase the correlation. Additionally, there may be three-way interactions between sexual selection, natural selection, and genetic drift. Therefore, future models that incorporate direct selection at both male trait and female preference loci in finite populations may be needed.

## Data Availability Statement

The Mathematica Notebook used for the perturbation analysis and the simulation code written in R is available at https://doi.org/10.5281/zenodo.14172252.

## Appendix

Let the vector *z* = (δ*t*, δ*p*, δ*D*) represent the deviations in allele frequencies and linkage disequilibrium (LD) from the deterministic trajectory in infinitely large populations. Let *z*^∗^ = *f*(*z*) be the recursion of *z*, which is described in the Method Section of the main text. Following Barton & Otto (2005), we assume that the deviations *z* remain small around 0, so the deviations in the next generation, *z*^∗^, can be expanded in a Taylor series as

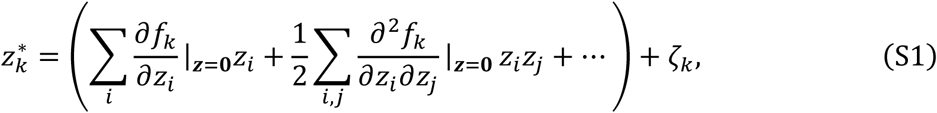

where *i*, *j*, *k* ∈ {*t*, *p*, *D*} refer to a dimension of *z*. The random variable, ζ_*k*_, is the perturbation due to sampling error after one generation (i.e., genetic drift). The first moments of these perturbations are 𝔼[ζ_t_ ] = 𝔼[ζ_p_] = 0, and 𝔼[ζ] = −D/2N. The second moments of the perturbations are

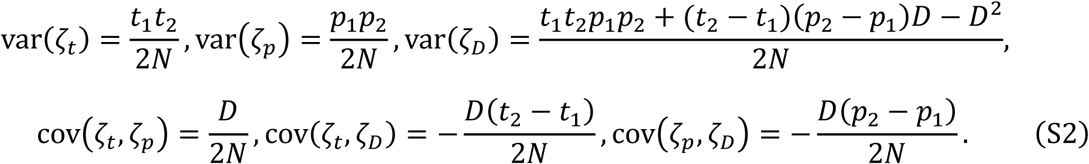

Based on Equation (S1), the expectation of the first and second moments of the deviations are

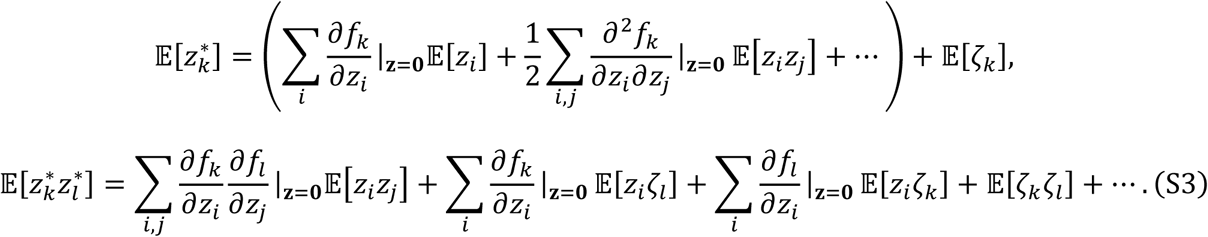

Assuming that deviations remain at the order of *O*(*N*^−1^) and ignoring *O*(*N*^−2^) terms, the first moment of the deviations are presented in Equation (1) in the main text. Dropping partial derivatives that are 0, the second moments of the deviations are

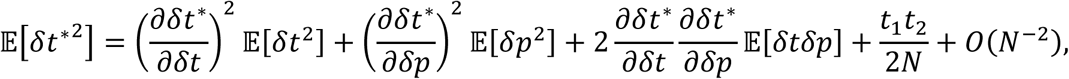

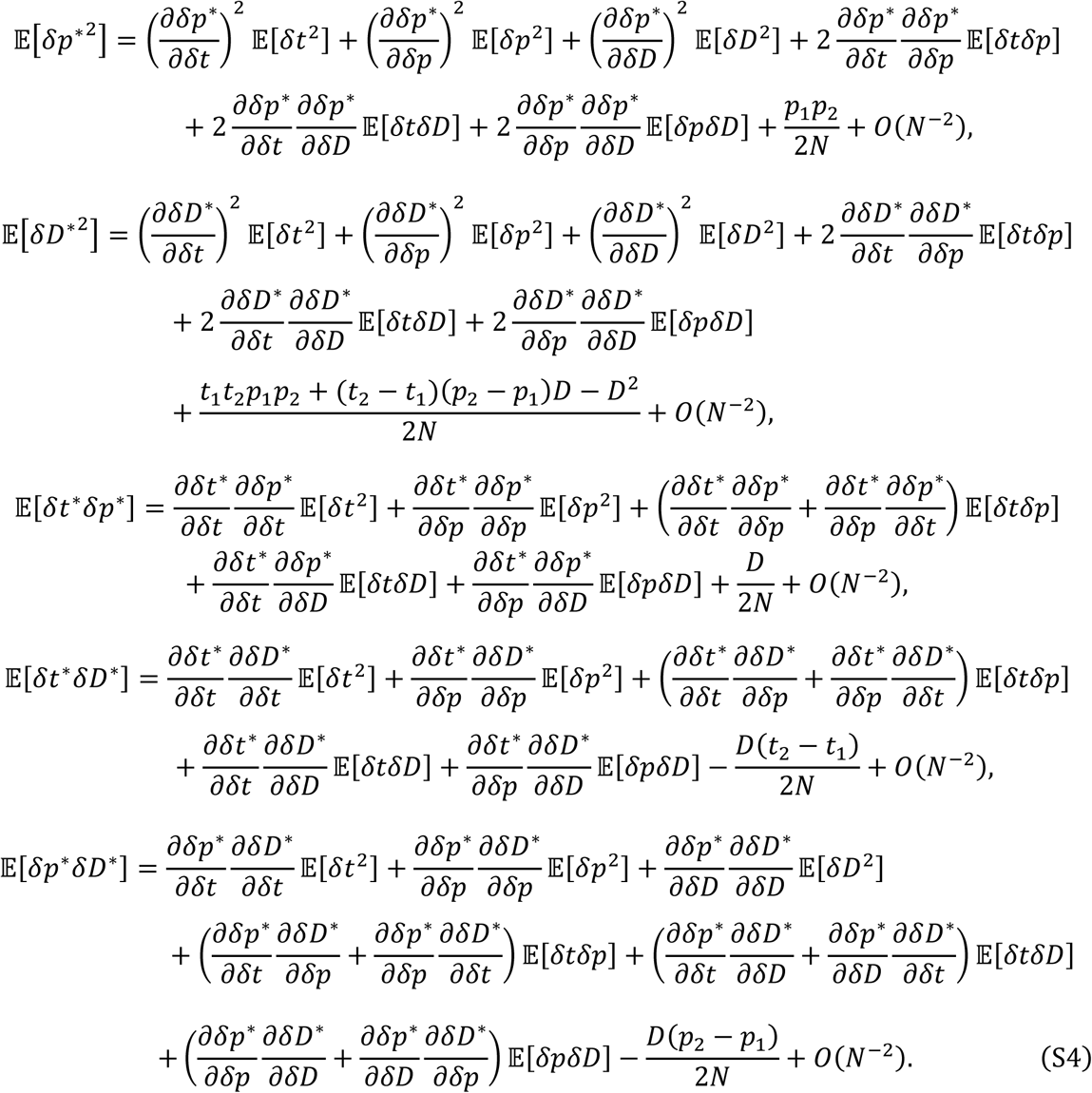

Note that recursions of terms involving the deviation in LD, δ*D*, depend on the deviations in allele frequencies (i.e., 𝔼[δ*t*], 𝔼[δ*t*^2^], 𝔼[δ*p*] and 𝔼[δ*p*^2^]). The expressions of the partial derivatives are complicated and are presented in the Mathematica Notebook.

## Notes

https://doi.org/10.5281/zenodo.14172252

### Competing Interest Statement

The authors have declared no competing interest.

