## Supplementary Figures for "When sexual selection meets genetic drift: the coevolution of male traits and female preferences in finite populations"

Kuangyi Xu[1]*

[1] Department of Ecology and Evolutionary Biology, University of Toronto, 25 Willcocks St, Toronto, ON, Canada M5S 3B2

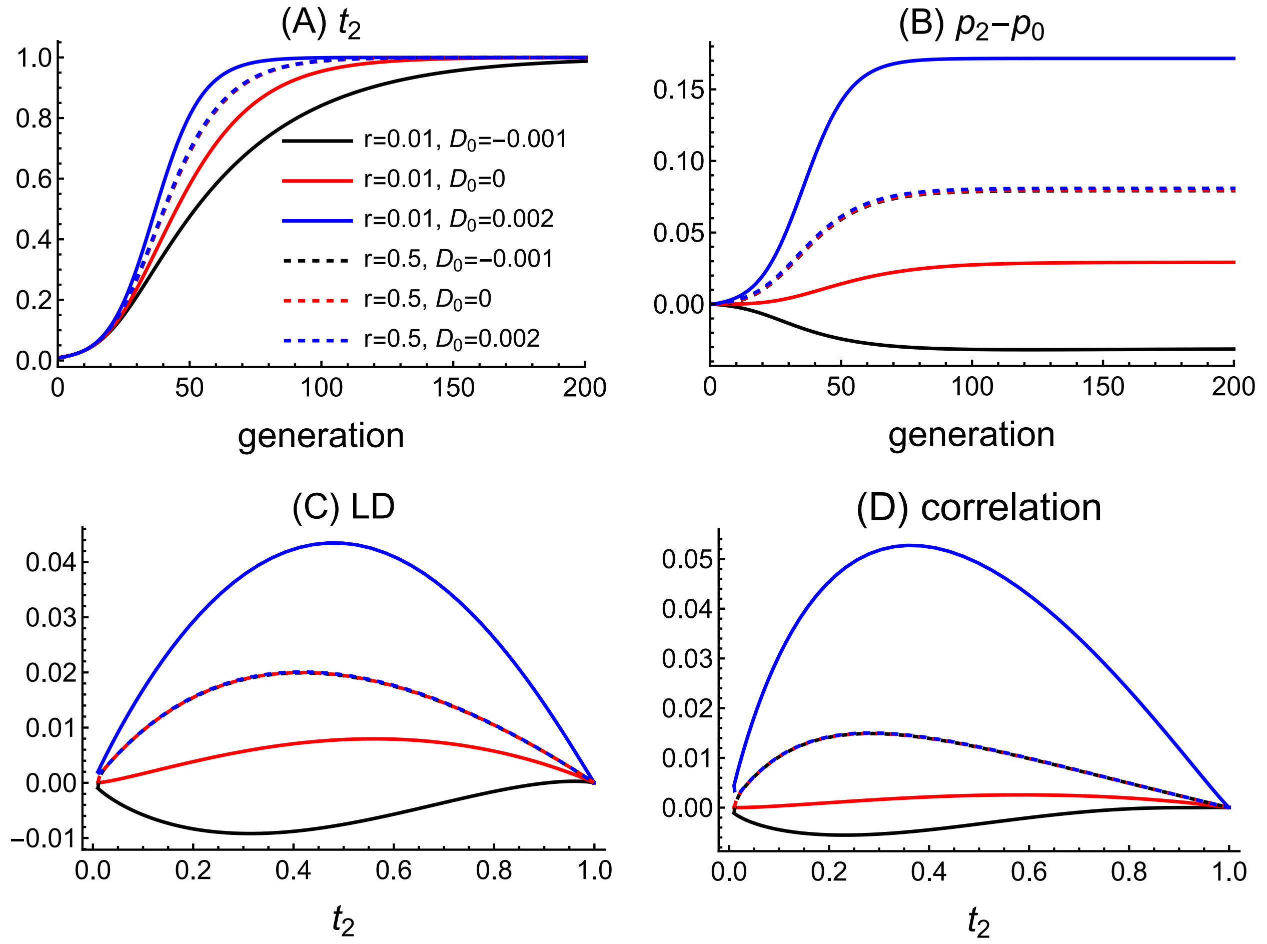


**Figure S1**. Effects of the initial LD between male trait and female preference alleles on the coevolution of the male trait and female preference in infinitely large populations. Solid and dashed lines show results under low and high recombination rates, respectively. Since LD depends on the genetic variation at both loci, to eliminate the impact of different genetic variation at the preference locus, we also plot the correlation coefficient of the male trait and preference alleles in panel (D). Other parameters used are $s = 0$, $a = 4$, $t_0 = 0.01$, $p_0 = 0.1$.

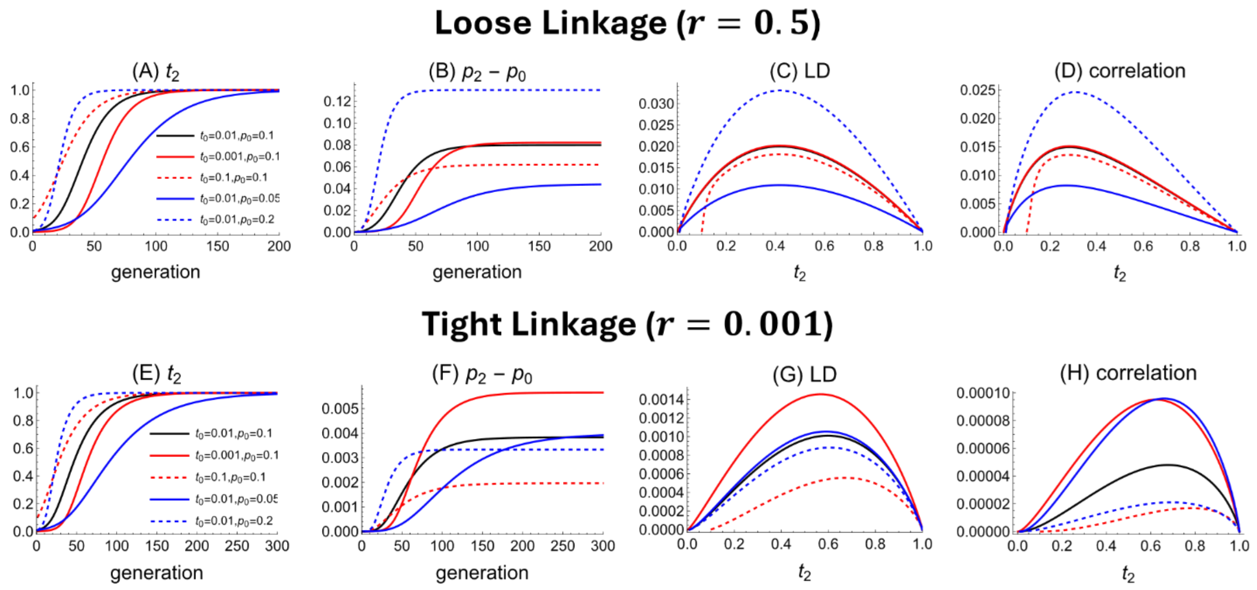


**Figure S2**. Effects of the initial frequency of the male trait allele ($t_0$) and the preference allele ($p_0$) on the coevolution of the male trait and female preference in infinitely large populations. Upper and lower panels show results under loose and tight linkage between male trait and preference loci, respectively. In the last two columns, the results are plotted against the male trait frequency rather than generations to account for different rates of evolution. Panels (D) and (H) correlation coefficient between trait and preference alleles to eliminate different genetic variation at the preference locus under different $p_0$. Unless otherwise specified, parameters used are $s = 0$, $a = 4$, $r = 0.5$, $t_0 = 0.01$, $p_0 = 0.1$, $D_0 = 0$.

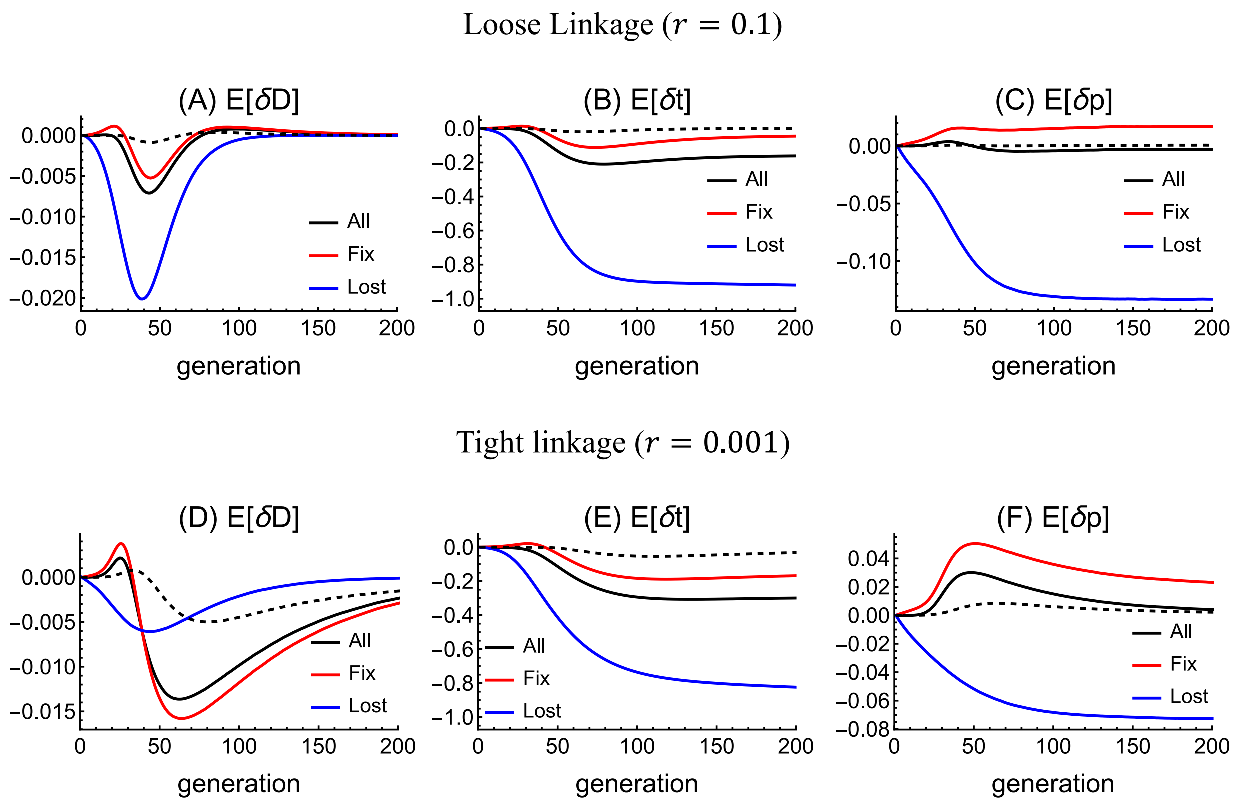


**Figure S3.** Expected deviations from the deterministic trajectory of $t_2$, $p_2$ and LD when the population size is small so that the perturbation approximation fails. Results are calculated from simulations. Solid lines depict the expected deviations in populations with $2N = 1000$, where the male trait allele $T_2$ may get lost. Black line is the deviation averaged across all replicates. Red and blue lines represent deviations averaged across replicates in which allele $T_2$ becomes fixed and lost, respectively. Dashed line show results at $2N = 10000$, in which case $T_2$ will always fix. Other parameters used are $a = 4, t_0 = 0.01, p_0 = 0.1, D_0 = 0$.

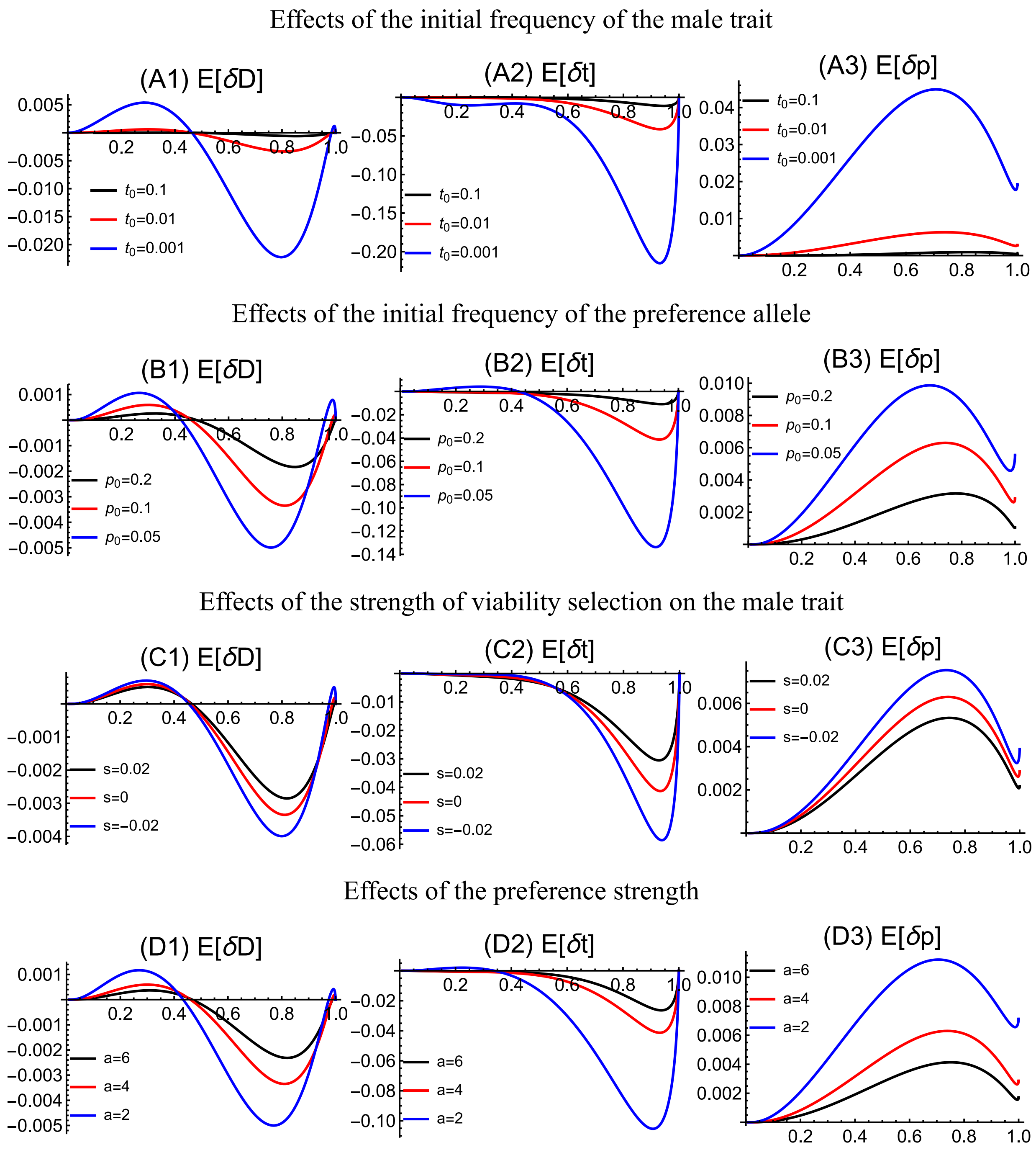


**Figure S4.** Factors that make the male trait $T_2$ sweep slower will amplify the magnitude of deviations. First row: influence of the initial frequency of the male trait $T_2$. Second row: influences of the initial frequency of the preference allele $P_2$. Third row; influences of selection coefficient of the male trait $T_2$. Fourth row: influences of the preference strength $a$. Unless otherwise specified, parameters are $2N = 10000, a = 4, s = 0, r = 0.01, t_0 = 0.01, p_0 = 0.1$.

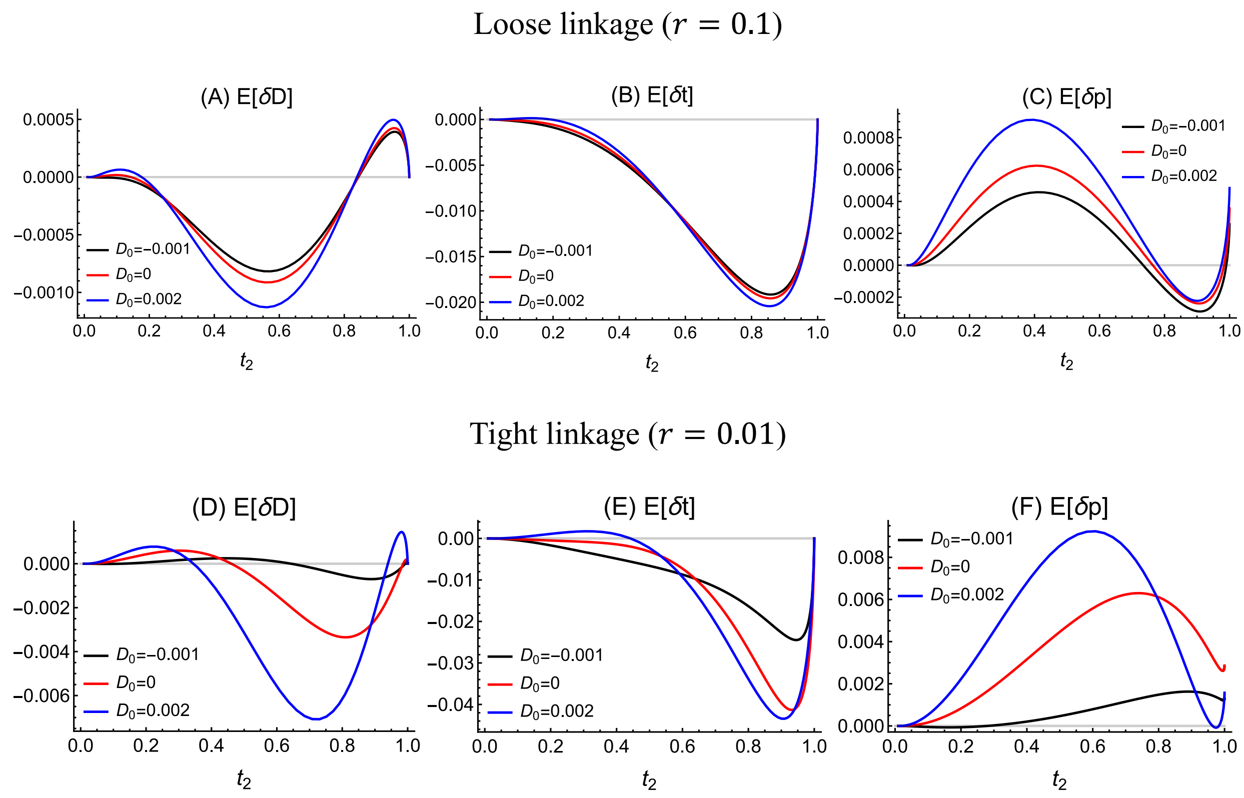


**Figure S5.** Impacts of the initial LD between male trait and female preference alleles on the deviations in LD and allele frequencies. Upper and bottom panels show results under loose and tight linkage, respectively. Other parameters are $2N = 10000$, $a = 4$, $s = 0$, $t_0 = 0.01$, $p_0 = 0.1$.

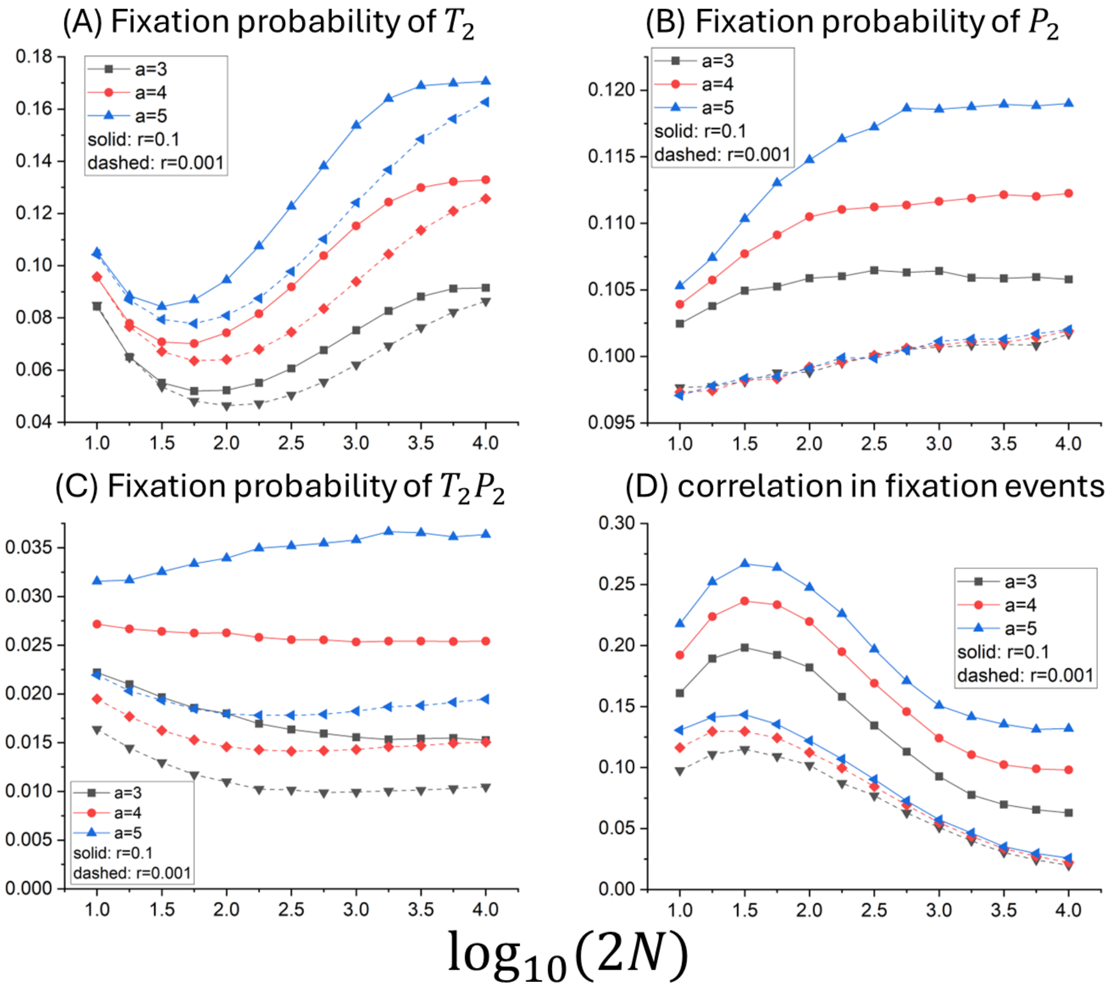


**Figure S6.** Impacts of the preference strength $a$ and population size on the fixation of a single male trait mutant and female preference allele. Parameter used are $p_0 = 0.1, s = 0$.

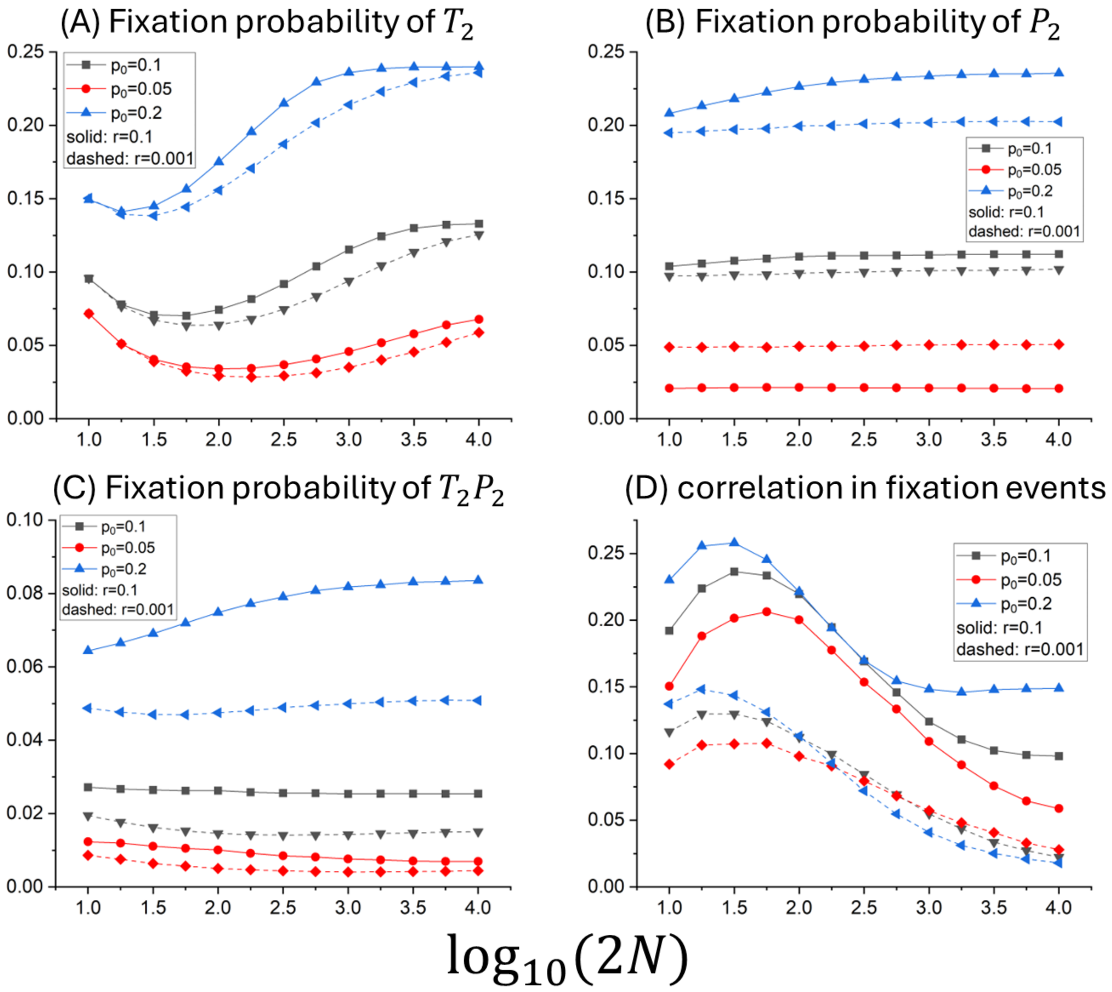


**Figure S7.** Impacts of the initial frequency of the preference allele ($p_0$) and population size on the fixation of a single male trait mutant and female preference allele. Other parameters are $a = 4, s = 0$.

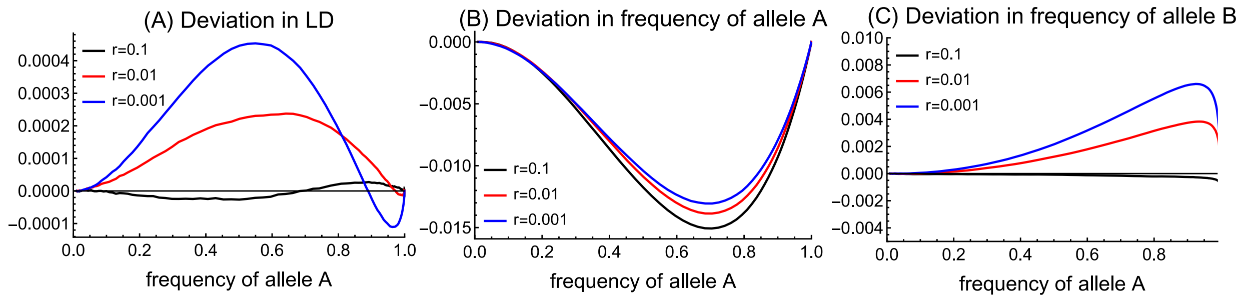


**Figure S8.** Linkage disequilibrium (LD) during the sweep of two beneficial alleles $A$ and $B$ under natural selection in a haploid population with non-overlapping generation. Different lines show results under different recombination rates between alleles. The selection coefficient $A$ is fixed to be $s_A = 0.1$, and the selection coefficient of allele $B$ is assumed to depend on the LD between the two alleles as $s_B = 0.2D$. There is positive epistasis, and the relative fitness of haplotype $AB$ is $1 + s_A + s_B + \epsilon$, with $\epsilon = 0.01$. Results are obtained from numerical simulations by averaging 100000 replicates. The initial frequencies of allele $A$ and $B$ are 0.01 and 0.05, respectively, with no initial LD.
